# Characterizing stochastic cell cycle dynamics in exponential growth

**DOI:** 10.1101/2021.08.24.457569

**Authors:** Dean Huang, Teresa Lo, Houra Merrikh, Paul A. Wiggins

**Affiliations:** Department of Physics, University of Washington, Seattle, WA, 98195; Department of Biochemistry, Vanderbilt University, Nashville, TN, USA; Department of Pathology, Microbiology, and Immunology, Vanderbilt University Medical Center, Nashville, TN, USA; Department of Bioengineering, University of Washington, Seattle, WA, 98195; Department of Microbiology, University of Washington, Seattle, WA, 98195

## Abstract

Two powerful and complementary experimental approaches are commonly used to study the cell cycle and cell biology: One class of experiments characterizes the statistics (or demographics) of an unsynchronized exponentially-growing population, while the other captures cell cycle dynamics, either by time-lapse imaging of full cell cycles or in bulk experiments on synchronized populations. In this paper, we study the subtle relationship between observations in these two distinct experimental approaches. We begin with an existing model: a single-cell deterministic description of cell cycle dynamics where cell states (*i.e*. periods or phases) have precise lifetimes. We then generalize this description to a stochastic model in which the states have stochastic lifetimes, as described by arbitrary probability distribution functions. Our analyses of the demographics of an exponential culture reveal a simple and exact correspondence between the deterministic and stochastic models: The corresponding state lifetimes in the deterministic model are equal to the exponential mean of the lifetimes in the stochastic model. An important implication is therefore that the demographics of an exponential culture will be well-fit by a deterministic model even if the state timing is stochastic. Although we explore the implications of the models in the context of the *Escherichia coli* cell cycle, we expect both the models as well as the significance of the exponential-mean lifetimes to find many applications in the quantitative analysis of cell cycle dynamics in other biological systems.

## I. INTRODUCTION

Methods to quantitatively characterize cell cycle dynamics have expanded dramatically [1] since the pioneering model of the *Escherichia coli* cell cycle described by Cooper and Helmstetter [2]. Their initial work represented the cell cycle as a deterministic process in which each step was precisely timed. Although these assumptions were almost certainly viewed as a matter of mathematical convenience, some later readers have interpreted the experimental success of this model as evidence that *all* the assumptions made to derive the model are supported by experimental evidence [3]. Some later authors have relaxed some of these assumptions and found that the predictions are in fact robust to the model details [3], but none have yet reanalyzed these dynamics in the context of the significant level of stochasticity observed in cell cycle timing (*e.g*. [4,5]).

One fundamental difficulty with reconciling the quantitative analyses of the cell cycle is the existence of two distinct classes of experiments: In *unsynchronized approaches*, an exponential culture is analyzed and the number of cells at time *t* is used to generate statistics defined with respect to cell number [2]. Examples of this approach are snapshot imaging (*e.g*. [6]), flow cytometry (*e.g*. [7]), and many deep-sequencing based approaches (*e.g*. [8]). We contrast these with *synchronized approaches* in which cells of a known state in the cell cycle progression are analyzed. Examples of this approach are the use of any of the previously described methods on cells which are first synchronized using a baby machine (*e.g*. [9]). Time-lapse imaging of full cell cycles (*e.g*. [10]), including the use of devices like the mother machine (*e.g*. [4]), can also be used to generate data for synchronized analyses. Although it might naïvely seem that averaging with respect to these two population ensembles are equivalent, they are not.

To demonstrate the subtlety of interpreting the data from an exponential culture, consider the probability of observing the Z ring, the ring-shaped protein complex that forms in *E. coli* at midcell and drives the process of septation (or cytokinesis) [11]. If the cell cycle has duration *T* and the Z ring has lifetime *δτ_Z_*, one might naïvely assume the probability of observing the Z ring is:

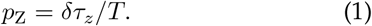

See Fig. 1A. Although this is true in the synchronized population, in an exponential culture the probability is 30% lower as a direct consequence of the relative abundance of cells by age [12]. Why? The number of new-born cells is twice the abundance of cells at the end of the cell cycle when the Z ring forms. Although this seems like a trivial book-keeping annoyance, when we consider the stochastic model, this effect has consequential implications for timing throughout the cell cycle, including on the growth rate.

**FIG. 1.**
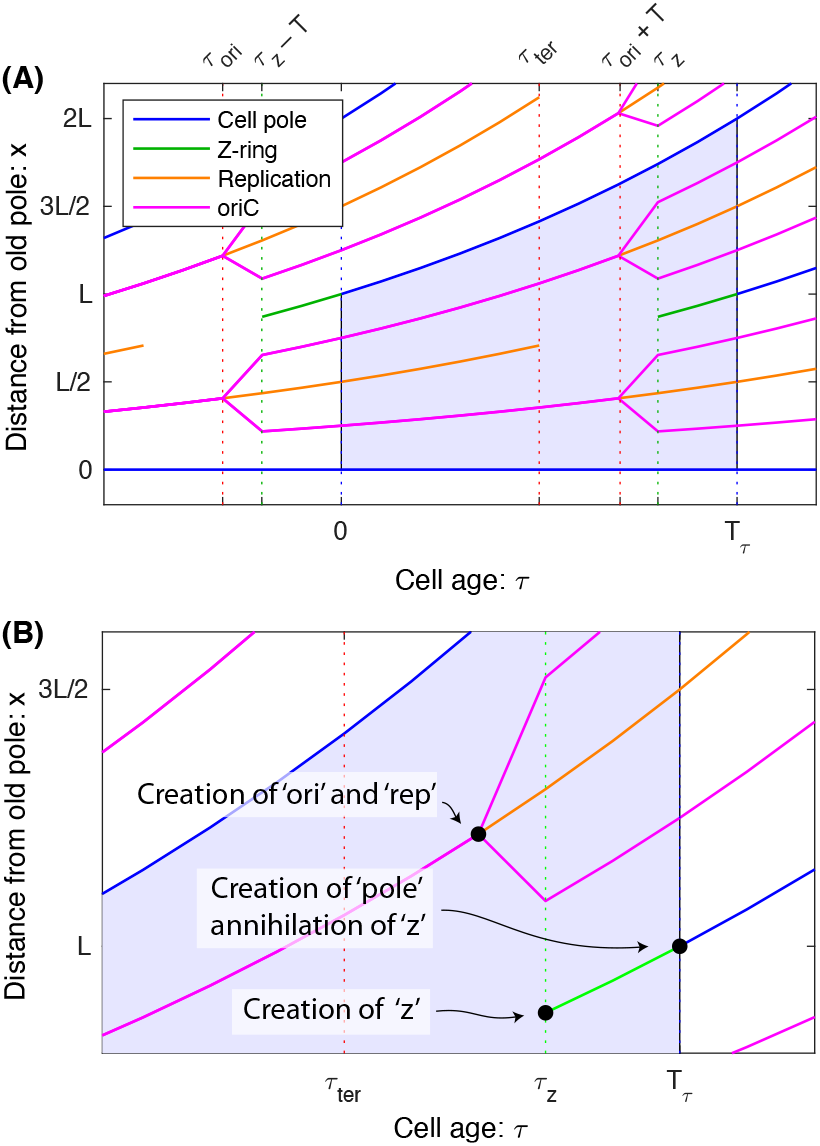
Panel A: Schematic for positioning and timing of events during the *E. coli* cell cycle. In the deterministic model, the duration of the cell cycle as well as the timing of all events in the cell cycle are precise (*i.e*. deterministic). Furthermore, we shall assume the positioning of all complexes and the length of the cell are all deterministic as well. One complete cell cycle is shaded in light blue. We have also annotated the positioning and timing of a number of cell cycle events: (i) Cell poles are annotated in blue and appear as the consequence of septation. (ii) The replication is annotated in orange, and starts when the origin (*ori*C) is replicated and ends when the terminus is replicated. The origin is replicated before the start of the cell cycle. (iii) The creation of new origins and their movement from the quarter-cell positions to the eighth-cell positions after replication is shown in magenta. (iv) The Z ring assembly, which drives septation at midcell, and then Z ring disassembly are annotated in green. **Panel B: Creation and annihilation.** The statistics of different types of quantities will have different statistical properties. *Transient quantities* undergo both creation and annihilation events. The Z ring and replisome are natural examples of transients since they assemble and then disassemble. In contrast, both cell poles and genetic loci are *perpetual quantities* that only undergo creation but never annihilation.

In Sec. IIA, we will first revisit the existing *deterministic model*, where all events in the cell cycle are precisely timed. In this model, we will represent the fundamental state of the cell as an age *τ* and compute the statistics of cell age in an exponential culture. To make contact with observables, we then apply these results to describe the demographics of the *E. coli* cell cycle in Sec. IIB. In Sec. IIC, we consider the *stochastic model*, where the cell cycle is represented as discrete sequential states *j* = 1…*m*, each with a stochastic lifetime *τ_δj_*. This model is also analytically tractable and we can compute expressions for all the same statistics as the deterministic model. However, the relation between the deterministic and stochastic model statistics is at this point opaque. In Sec. IID, we define an *exponential mean*, which is a mean biased toward younger cells that are overabundant in exponential culture. In Sec. IIE, we demonstrate that the predictions of the stochastic and deterministic models are in fact identical if the deterministic state ages *τ_j_* are equal to the exponential-mean stochastic state ages 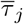. Finally, in Sec. IIE, we consider a number of simple biological examples to underline both the mathematical behavior of the exponential mean as well as its biological implications. In the interest of brevity, we will discuss experimental support for this model elsewhere [13].

## II. RESULTS

In this section, we will derive the expressions for a large number of statistics relevant for describing an exponential culture. We will first derive expressions for the statistics in the deterministic model and then the stochastic model. In Tab. I, we provide a summary of the notation.

**TABLE I.** A summary of the model notation. Note that since the deterministic model is described in terms of a continuous state, the age *τ*, the number of cells in that state is represented as a *number density n*; whereas in the stochastic model, the cell state is represented by an integer *j* and, therefore, the number of cells in that state is represented by a *number N_j_*. The symbol ~ means that a random variable is *distributed like*.

| Variable | Meaning |
| --- | --- |
| $t$ | Experimental time |
| $N(t)$ | Total number of cells at time $t$. |
| $N_{\text{obj}}(t)$ | Number of object 'obj'. |
| $k$ | Culture growth rate |
| $T$ | Culture mass doubling time |
| **Deterministic model** |  |
| $\tau$ | Cell age $0 \leq \tau \leq T_\tau$ |
| $T_\tau$ | Deterministic duration of the cell cycle |
| $N_\tau^+(t)$ | Cumulative number of cells that have had age $\tau$ |
| $n_\tau(t)$ | Number density with respect to cell age $\tau$ |
| **Stochastic model** |  |
| $\tau_{\delta j} \sim p_{\delta j}(\cdot)$ | Stochastic lifetime of state $j$ |
| $\tau_j \sim p_j(\cdot)$ | Stochastic cumulative lifetime of states $\leq j$ |
| $T_\tau \sim p(\cdot)$ | Stochastic duration of the cell cycle |
| $N_j(t)$ | Number of cells in state $j$ |
| $N_j^+(t)$ | Cumulative number of cells that have been in state $j$ |

### A. Deterministic model

In the deterministic model, we will consider cells that are born with age *τ* = 0 and divide deterministically at age *τ* = *T_τ_*. By *cell age τ*, we mean a continuous cell state variable representing cell cycle progression, not *aging* in the context of reduced cell fitness over time [14].

#### 1. Definition of the deterministic model

Since cell age *τ* is continuous, we define a number density *n_τ_*(*t*) of cells with age *τ* at time *t*. Age *τ* is defined on the interval [0, *T_τ_*] with *τ* = 0 corresponding to cell birth and *T_τ_* corresponding to cell division. We define the *doubling time T* as the mass doubling time in an exponential culture, which we distinguish from the cell cycle duration *T_τ_*. Let the creation number, 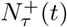, be the cumulative number of cells that have had age *τ* and the annihilation number, 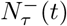, be the cumulative number of cells past *τ* age. To describe this model formally, we write rate equations for the number of cells with an age between *τ* and *τ* + *δT*:

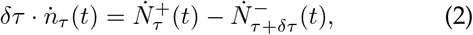

where *Ȧ* ≡ *∂_t_A*. In the deterministic model, the number of cells entering and exiting the age-*τ* state are:

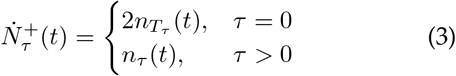

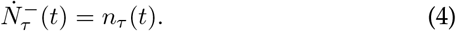

Substituting Eqs. 3-4 into Eq. 2 gives a single rate equation in terms of the number density *n*:

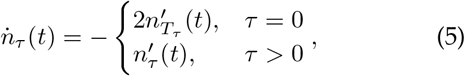

where *A*′ ≡ *∂_*τ*_A* and the *τ* = 0 case corresponds to the cell division event and the creation of two daughters age *τ* = 0. Eq. 5 completely describes the cell cycle dynamics in the deterministic model.

#### 2. Solution to the deterministic model

In steady-state growth, we can assume the total number of cells is:

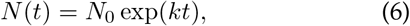

where *k* is the growth rate that is determined by solving the rate equation (Eq. 5). It will often be convenient to rewrite the equations in terms of the doubling time:

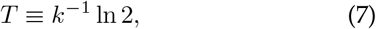

rather than the growth rate *k*. Eq. 5 evaluated at *τ* = 0 gives a consistency condition between doubling time *T* and the duration of the cell cycle *T_τ_*:

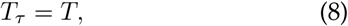

which is to say that the doubling time is equal to the duration of the cell cycle, as one would naïvely expect. In steady-state growth, one can compute the number density of cells, which is:

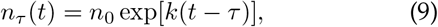

where *n_τ_* is the density with respect to cell age *τ* and *n*_0_ is a constant determined by the initial cell number. The details of the derivation of the solution are given in the Appendix A1.

#### 3. Statistics of the deterministic model

The solution of Eq. 5 can be used to compute the probability (PDF) and cumulative (CDF) distribution functions with respect to cell age:

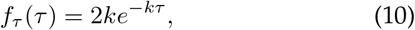

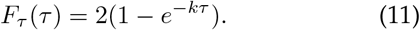

Eq. 10 implies that in an exponential culture, there is an enrichment of young cells which decays exponentially with age *τ*. See Fig. 2A.

**FIG. 2.**
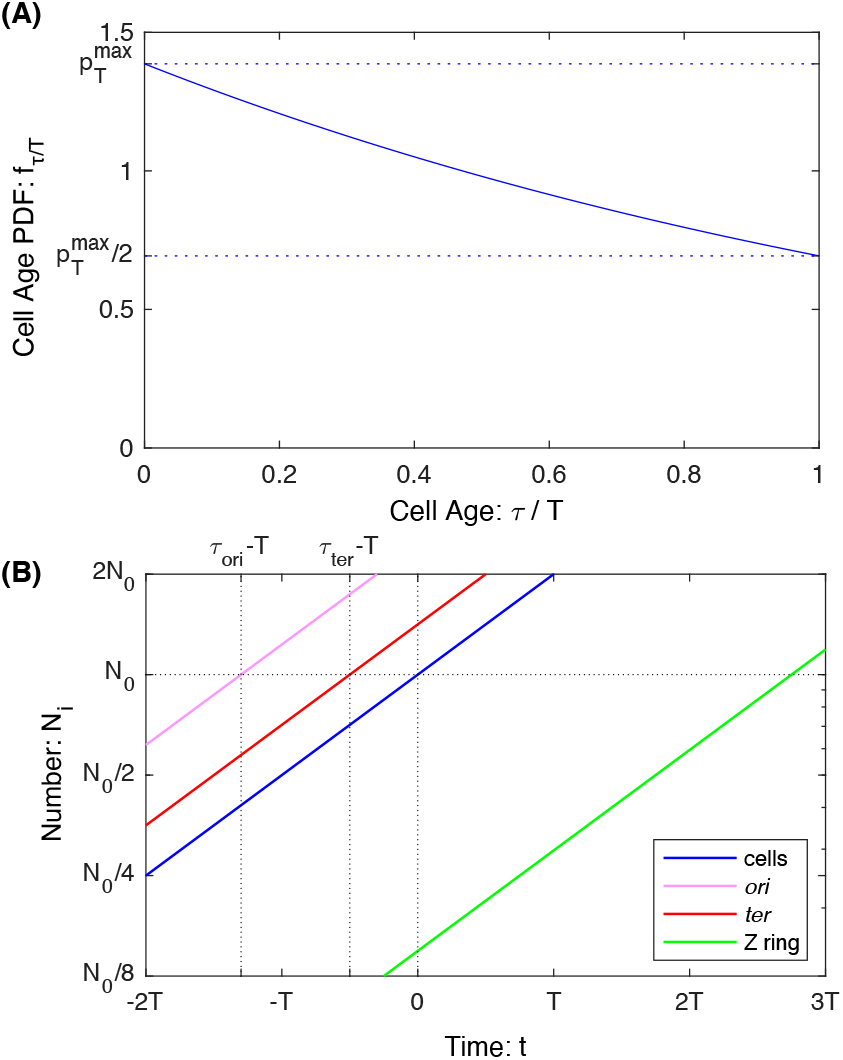
Panel A: Cell Age PDF. In an exponential culture, there is an enrichment in young cells relative to old cells. The relative number of cells decays exponentially with cell age *τ*. **Panel B: Numbers in an exponential culture.** The numbers of all quantities grow exponentially with the same growth rate. For perpetual quantities (*e.g*. cell number, cell poles, DNA loci) the relative timing of the creation of a quantity can be inferred by the temporal offset of the *N_X_*(*t*) curve relative to the cell number curve *N*(*t*). In contrast, transient quantities, like the number of Z rings, also grow exponentially, but their offset cannot directly be interpreted as a time.

Note that the canonical observable in an exponential culture is number as a function of time rather than abundances relative to the total number of cells *N*(*t*). However, we shall write each expression as the prefactor of *N*(*t*) and therefore the prefactor can be interpreted as the abundance relative to cell number *N*(*t*).

The creation number is:

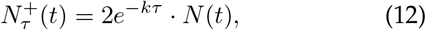

which is the cumulative number of cells to go through age *τ*. The number of cells younger than age *τ* is:

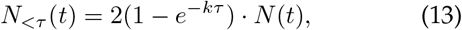

and the number of cells older than age *τ* is:

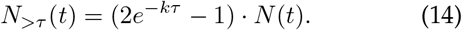

Finally, the number of cells in a state defined by the age range *τ*_1_ < *τ* < *τ*_2_ is:

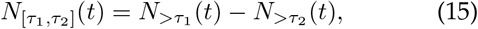

as derived in Appendix A1.

### B. Application to cell cycle dynamics

In this section, we demonstrate how to apply these results in the context of the *E. coli* cell cycle dynamics shown schematically in Fig. 1A. These formulae can be applied either to predict the numbers from the known replication timing or to infer timing from the observed numbers in an exponential culture.

#### 1. Z ring

First consider the Z ring: The Z ring is an example of a transient complex, therefore we need to use *N* (as opposed to *N*^+^). Furthermore, it assembles at *τ* = *τ*_z_ and disassembles at the end of the cell cycle. The number of Z rings is therefore equal to the number of cells older than *τ_Z_*:

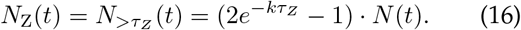

It is interesting to consider the limit as *δτ_Z_* ≡ *T_τ_* – *τ*_Z_ is small relative to the cell cycle duration *T_τ_* in order to compare this to our intuitive guess (Eq. 1):

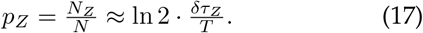

Since ln 2 ≈ 0.69, this is roughly 30% smaller than our naïve estimate due to depletion of older cells in an exponential culture (Fig. 2A).

#### 2. Cell poles

The number of cell poles is obviously twice the number of cells, but it is useful to consider this example more formally. Unlike the Z ring which is transient, the poles are perpetual: Once the state is created, it is never annihilated (Fig. 1B). In this context, we can use the creation number *N*^+^. Note that we are immediately presented with a conundrum: Are two poles formed at the end of the cell cycle (*τ* = *T*) or is one pole created at birth (*τ* = 0)? Both approaches give the same number:

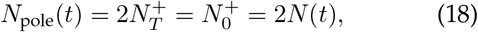

which is twice the number of cells, just as one intuitively expects.

#### 3. DNA loci

Now we consider the number of a particular genetic locus *ℓ*. First consider the slow-growth limit where both initiation and termination occur within the current cell cycle [2]. Assume the locus of interest is replicated at time *τ_ℓ_*. The number of copies per cell is one before replication and two after replication. We can therefore write:

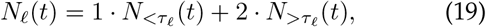

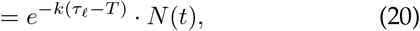

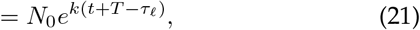

in agreement with previous results [15,16]. Unlike transient quantities, *e.g*. the number of Z rings, the form of Eq. 21 implies that the number of genetic loci can be understood as a temporal shift of *N*(*t*) by *T_τ_* – *τ_ℓ_* to shorter times, as illustrated in Fig. 2B.

The cancelation between the non-exponential terms between *N*_>*τ_ℓ_*_ and *N*_<*τ_ℓ_*_ in Eq. 20 may seem incidental, but from another perspective it is intuitive: The mathematical reason for the non-exponential terms in the prefactors of Eqs. 13-14 is *annihilation, i.e*. the reduction in the number of cells of a particular age *τ* due to aging. DNA loci correspond to a perpetual state: Once a locus state is created (*i.e*. replicated) it does not annihilate (*i.e. transition* into another *state*). To compute the number of genetic loci, we can therefore use the creation number *N*^+^ formula (as opposed to *N* which is reduced by annihilation). This more direct approach yields the same result as Eq.20:

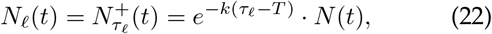

but is applicable for fast growth where replication initiates before the cell cycle begins (i.e. *τ_ℓ_* < 0), as illustrated in the cell cycle schematic in Fig. 2A.

#### 4. B, C, and D period

The B period is defined as the period between birth and replication initiation. The C period is defined as the cell cycle period during which replication occurs: *i.e*. after replication initiation and before termination [2]. The D period is defined as the period between replication termination and cell division [2]. The relation between the durations of these periods and the locus number (Eq. 22) are:

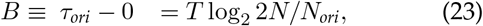

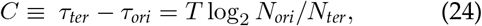

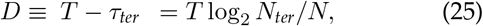

where *N, N_ori_*, and *N*_*ter*_ are the number of cells, origins, and termini in the exponential culture (not per cell), which has previously been reported [2,3]. Note that if replication initiates before the start of the cell cycle, *B* = *τ*_*ori*_ < 0.

#### 5. Replication

Finally, let us consider the replisomes and the replication process itself. Like the Z ring, this is a transient state. However, there is a significant subtlety here: Do we count (i) replicating cells, (ii) individual replication processes consisting of replisome-pairs, or (iii) individual replisomes?

First let us consider the number of replisome-pairs. Since the replication process can span the overlap between two successive cell cycles, it is most convenient to use differences in the creation number *N*^+^. In fact, we can express it concisely in terms of *ori*C and *ter*:

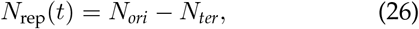

and the number of individual replisomes will be twice the number of pairs. *N_ori_* and *N_ter_* are computed using Eq. 22.

For the number of replicating cells, we consider three different cases. First consider a case where the replication cycle is internal to the cell cycle. In this case, we have:

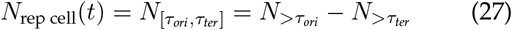

which can be evaluated using Eq. 15. If the replication process overlaps by a single cell cycle but replication rounds do not overlap:

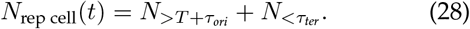

Finally, if the rounds of replication overlap:

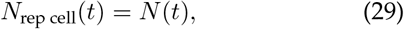

and all cells are replicating in the deterministic model.

### C. Stochastic model

An important complication of a more realistic model for cell cycle dynamics is stochasticity (*i.e*. randomness) in the lifetime of the states of the cell cycle. We will represent this stochasticity by dividing the cell cycle into *m* discrete states through which the cell must transition sequentially. This model is shown schematically in Fig.3. The lifetime of each state *τ_δj_* will be described by an arbitrary lifetime PDF for the *j*th state *τ_δj_* ~ *p_δj_*(·).

**FIG. 3.**
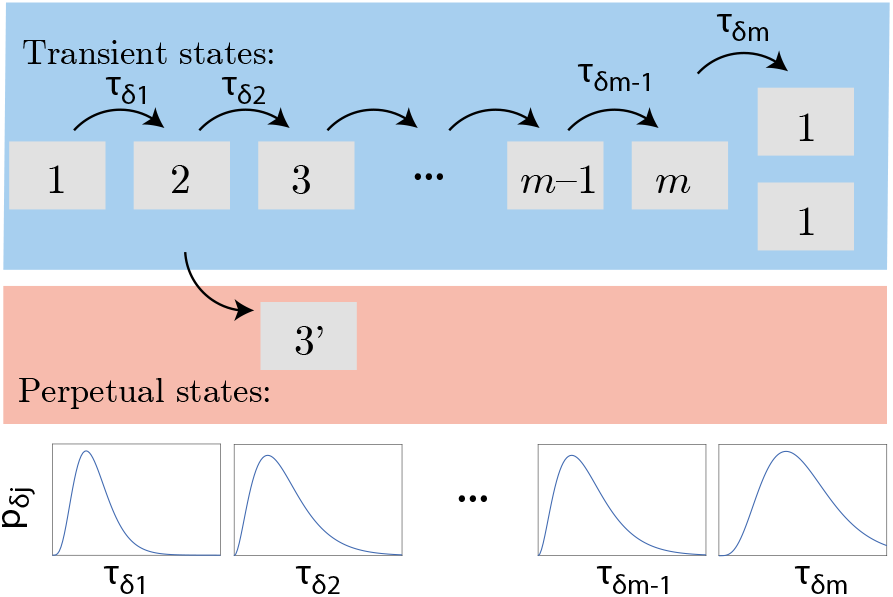
Schematic of the stochastic model of the cell cycle. We represent the cell cycle as a series of sequential states *j* = 1…*m*. After the *m*th state, two daughter cells in state 1 are born. The PDF of state lifetimes, *p_δj_*, is distinct for each state *j*. Although most quantities of interest are transient, meaning that the states are *created* when the cells enter and then *annihilated* when the cells exit, we also consider perpetual quantities that are created but not annihilated (*e.g*. DNA loci).

#### 1. Definition of the stochastic model

Let *N_j_*(*t*) be the number of cells in state *j*, the creation number, 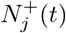, and the annihilation number, 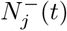, be the total number of cells to have arrived and departed from state *j*, respectively. The state dynamics is therefore described by the following rate equation:

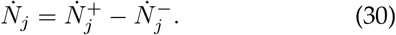

In this model, cells move sequentially through the *m* states before the final state (*j* = *m*) transitions to the initial state (*j* = 1) as two cells:

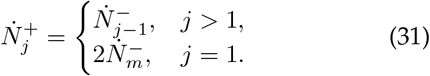

Each state *j* has a PDF of lifetimes *p*_*δj*_(*t*) and therefore the relation between state *j* arrivals and departures is given by:

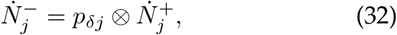

where ⊗ is the convolution:

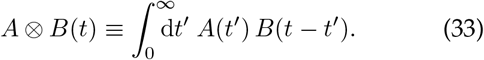

Eqs. 30-32 completely specify the stochastic model.

#### 2. Solution to the stochastic model

We will work in the steady-state growth limit as before. It is most convenient to work in terms of the Laplace transforms of the rate equations (Eqs.30-32). The Laplace transform is defined:

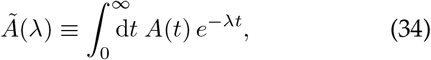

where the tilde denotes the transformation from time *t* to Laplace conjugate λ.

The transformed representation is convenient since the ordinary differential equations become algebraic equations in terms of the transform quantities and the convolutions become products of transforms (*e.g*. [17]).

Of particular importance in what follows is the relation between the PDF of lifetimes of individual state *j*, *p_δj_*(*t*), and the PDF of times taken to transition from birth through state *j*, *p_j_*(*t*):

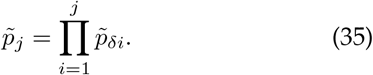

A detailed derivation of the solution to the rate equations (Eqs.30-32) is given in Appendix A2.

For steady-state exponential growth, the consistency condition that relates the growth rate *k* to the PDF of cell cycle durations *p*(*t*) ≡ *p_m_*(*t*) can be written concisely in terms of the Laplace transform:

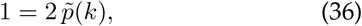

an equation that is well known [18,19]. This consistency condition is equivalent to Eq. 8 in the deterministic model, although the mathematical equivalence between these two relations is opaque for the moment.

#### 3. The statistics of the stochastic model

In an exponential culture, the cell numbers are given by the expressions:

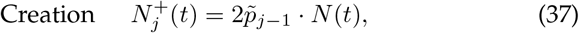

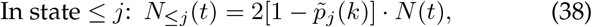

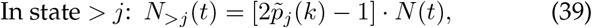

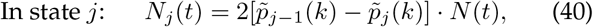

which have a similar structure to the dynamics of the deterministic model (Eqs. 12-15), but are dependent on the Laplace transforms of the state lifetime PDFs. Intuitively, these Laplace transforms give rise to an effective mean time.

### D. The exponential mean

To understand the biological significance of the Laplace transform of the lifetime PDF, consider the generalized f-mean (or Kolmogorov mean) where the random variable *t* is first transformed by function *g*, an arithmetic mean is performed, and then the inverse function is applied to generate a generalized expectation [20]:

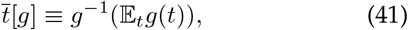

where 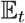 is the arithmetic expectation over random variable *t*. Both the harmonic mean and geometric mean are special cases of this more general formulation. The Laplace transform of the state-lifetime PDF can be reinterpreted as the expectation of *g*(*t*) = exp(–*kt*) and therefore we can generate the f-mean:

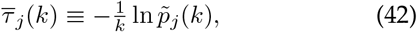

which can be understood as the exponential-mean of the lifetime of state *j*.

Before returning to our model, we will explore the behavior of the exponential mean. Consider the special case of a distribution that is very narrow relative to the growth rate. In this case:

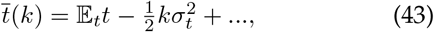

where the exponential mean is equal to the mean to the order of the variance 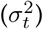 times the growth rate *k*. Short-lived states and states with small lifetime-variance will therefore have exponential means equal to the mean. More generally, the Jensen inequality always guarantees the exponential mean is less than or equal to the mean:

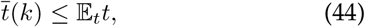

since the function *g*(*t*) is convex [21].

Finally, let us consider the consequences of a very wide distribution of lifetimes. Consider a state *j* in which fraction *∊* of cells arrest (*τ_j_* → ∞) while the remaining cells have exponential-mean lifetime 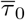. Using Eq. 42, it is straightforward to compute the exponential-mean lifetime:

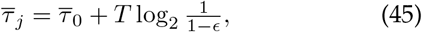

where the second term acts to extend the lifetime by a positive multiple of the doubling time *T*. Although the arrested cells do lengthen the exponential-mean lifetime, it remains finite. Eq. 45 is a useful approximation anytime some fraction of the cells have a lifetime much longer than the doubling time even if all lifetimes are finite.

### E. Model correspondence

To determine the differences between the deterministic and stochastic model, we eliminate the Laplace-transformed lifetime PDFs 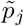 in favor of the exponential-mean lifetimes 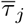 using their definition (Eq. 42). First consider the consistency condition for exponential growth (Eq. 36). The convolution theorem ensures that the exponential-mean times of successive states add to generate the wait time for transitioning from birth to state *j* (e.g. the natural logarithm of Eq. 35). Eq. 36 can now be rewritten as a relation between the cell cycle duration and the doubling time:

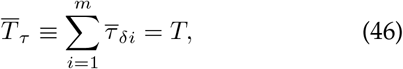

which is now intuitively equivalent to the consistency condition in the deterministic model (Eq. 8).

Now consider the expressions for state number in the stochastic model (Eqs. 37-39). When the deterministic cell age is set equal to the exponential-mean stochastic transition time 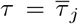 for the state *j*, the numbers are identical in the two models:

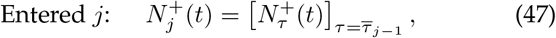

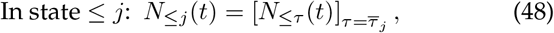

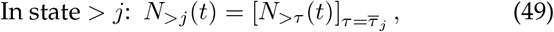

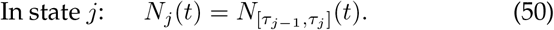

We therefore conclude that the statistics of the deterministic and stochastic models are identical in an exponential culture for models with the same growth rate *k*, once a correspondence has been established between states *j* and ages *τ*. In the deterministic model, state *j* corresponds to times 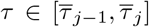 where 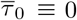 and 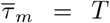. This correspondence is illustrated schematically in Fig. 4. Since we demonstrated a correspondence between the models, almost all the application discussed in Sec. IIB generalize by replacing the deterministic time *τ* with the corresponding exponential mean 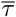 [22].

**FIG. 4.**
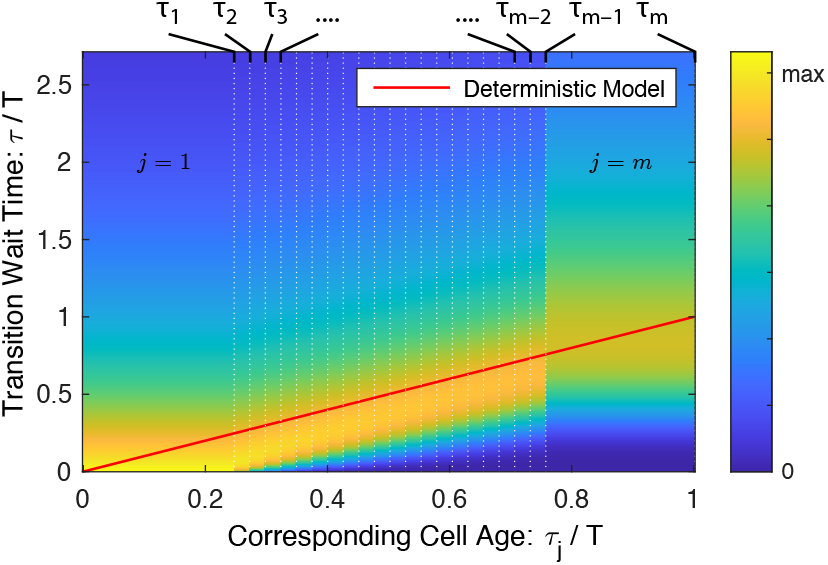
Model correspondence. The deterministic and stochastic models generate identical statistics in exponential growth once a suitable correspondence is defined between cell state *j* in the stochastic model and cell age *τ* in the deterministic model. State *j* in the stochastic model corresponds to age interval 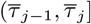 in the deterministic model, which is represented by the red line. In the stochastic model, the PDF of lifetimes as a function of state *j* is shown. Quantitatively, the age in the deterministic model tracks with the peak lifetime in the stochastic model.

### F. Implications for cell cycle phenomenology

To explore the nontrivial consequences of stochasticity in timing, consider an example motivated by replication conflicts [23]: By visualizing the replisome dynamics using single-molecule microscopy, we have recently reported that transcription leads to pervasive replisome instability [24]. To what extent should conflict-induced pauses in replication have been detectable in the classic analyses of unsynchronized cell populations?

Consider a simplified model in which an experiment probes the difference between the wildtype strain W and two mutant strains. The wildtype W grows with deterministic C period *C*_W_ and deterministic cell cycle duration *T*_W_. In mutant strain A(rrest), a small fraction e of cells arrest during replication (*i.e*. C period) and never complete the cell cycle, whereas non-arrested cells are identical to wildtype cells. In mutant strain S(low), the replication process is 1 + *∊*′ times slower, but the B and D periods are identical to wildtype. Using Eq. 45, one can compute the C period duration *C* and doubling time *T*. To lowest order in **∊**, the inferred cell cycle durations and C period are presented in Tab. II.

**TABLE II.**
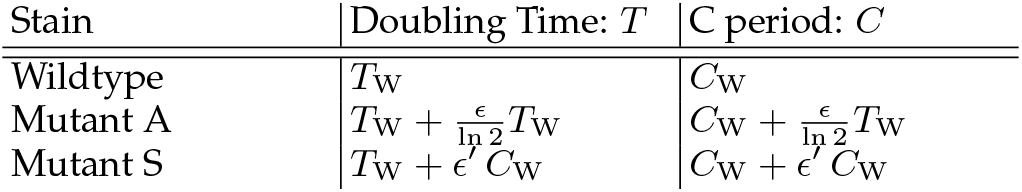
The effect of mutants on the doubling time *T* and C period duration *C* of an exponential culture.

At an intuitive level, one aspect of the prediction is easy to understand: In both mutants the C period is lengthened, as one might naïvely expect since this is the replication period of the cell cycle. Furthermore, the doubling time increases by the same amount as the C period increases. But there is an aspect of this prediction which is perhaps less intuitive: One might naïvely expect to observe a more dramatic consequence of replication arrest, like a large buildup of C period cells, but the consequences are indistinguishable from a slowdown in an exponential culture. In both mutants, there is a slight lengthening of the inferred C period, even though the slowdown is caused by replication arrest in the context of the A mutant. Although this prediction is not new in a qualitative sense, it concisely illustrates how the statistics of the exponential culture mask two mechanistically distinct phenomena.

The statistics of an exponential culture can also generate distinctions where seemingly none exist. Consider a more realistic model in which the duration of the D period is stochastic, has a non-zero width, and is *identical* for all three strains. The more rapid growth rate of the wildtype strain implies that its effective D period is shorter than for mutants cells:

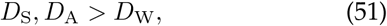

even though the distributions of the D period durations are identical in all three strains. (To understand how this occurs, see the second term on the RHS of Eq. 43.) In most cases this effect should be subtle, but for large changes in growth rate, these changes could be quite significant and can clearly complicate the interpretation of effective period lifetimes in an exponential culture.

## III. DISCUSSION

In this paper we provide a detailed analysis of both deterministic and stochastic models of the cell cycle. In Sec. IIA, we solved the deterministic model in which the cell-aging and division processes are precisely timed and determined the demographics (*i.e*. statistics) of an exponential culture. Given a set of observed demographics, Sec. IIB provides a detailed road map for how to infer cell cycle state timing in the context of the deterministic model. In Sec. IIC, we solved the more realistic stochastic model in which the lifetimes of sequential states are stochastic and again we determined the demographics of an exponential culture. By defining an exponential mean in Sec. IID, we demonstrated that the statistics of the two models were equivalent in Sec. IIE. The effective lifetime of states in the deterministic model is the exponential mean of the lifetimes in the stochastic model. That is to say that the exponential-mean lifetimes are the *sufficient statistics* of the model (*e.g*. [25]): Knowledge of only these lifetime statistics predicts the demographics of the exponential culture; therefore, inference on exponential-culture demographics infers only the exponential means, rather than the underlying lifetime distributions themselves. Finally in Sec. IIF, we discussed some of the limitations of the exponential-mean lifetimes in resolving the underlying biological mechanisms.

### A. Applicability of the stochastic model

Is the stochastic model sufficiently complex to capture all the relevant cell cycle phenomenology in *E. coli* and other bacterial systems? Like the deterministic model before it, the stochastic model is an idealized model that is simple enough to be tractable analytically, but complex enough to capture some important phenomenology. There are a number of shortcomings of this model but perhaps the most significant is that there is *no memory* beyond the cell state index *j*. As a consequence, it makes predictions at variance with some observed phenomenology: For instance, the stochastic model must predict that successive cell cycle durations are uncorrelated; however, these correlations are observed [4]. (We briefly consider the implications of a more general model in Appendix A3.) Another important limitation of the stochastic model is that cell divisions are symmetric, which is a good approximation in *E. coli*, but these types of stochastic models can easily be extended to the general asymmetric division case (*e.g*. [19]).

### B. On the applicability of the exponential mean

Although the definition of the exponential mean was motivated by the correspondence between the deterministic and stochastic models, it almost certainly has much greater applicability to other more complicated scenarios. For instance, our own numerical experiments using more complex models suggest that the relation between the effective lifetime of the states and the exponential-mean lifetime appears to be more robust than the assumptions of the stochastic model might imply. Since the key mechanism for generating bias toward short times is steady-state exponential growth, we expect the exponential mean of wait times to be the determinative statistic in more general models, as demonstrated in Appendix A3. As such, the exponential-mean lifetime could be a powerful observable to bridge timescales between single-cell and culture phenomenology in two different contexts: (i) in experiments probing cell cycle dynamics at the single-cell level and (ii) in complex numerical simulations that are too slow and too memory intensive to simulate in the long time limit [26].

We should note that although we believe our interpretation of the doubling time as an exponential mean (Eq. 46) is novel, it has already been appreciated in two important respects: (i) From a computational perspective, the Laplace-transform formulation (Eq. 36) of Eq. 46 has long been known [18]. (ii) From a qualitative perspective, biologists have long understood the consequences of the exponential-mean lifetime on cell growth rate: *I.e*. the doubling time *T* is “an average” of the cell cycle duration *T_τ_*; however, a small arrested subpopulation, for whom *T_τ_* → ∞, slows but does not stop growth. There is also physical precedent for this type of mean: Intriguingly, it emerges in context of non-equilibrium statistical mechanics [27] [28], although what connection this has to our cellular dynamics is opaque.

### C. On the significance of stochasticity

How does stochasticity affect biological function? Experimentally, we have long known that although the statistics of an exponentially growing population are well described by the deterministic model [2], nontrivial stochasticity in cell cycle timing is observed [4,5]. It is therefore tempting to conclude, based on the literature and perhaps even our own results, that stochasticity is either *small* or simply *does not* significantly affect biological function.

Our own conclusions are much more nuanced. Although our results guarantee that the deterministic model fits the exponential-culture demographic data just as well as the stochastic model, we have demonstrated that the stochasticity in timing is hidden in plain sight. The distribution of state lifetimes determine the exponential means. Therefore, the success of the deterministic model should not be interpreted as evidence against stochasticity or against its importance, but rather it indicates that only the exponential-mean state lifetimes are determinative parameters in the model for the demographics of an exponential culture.

Perhaps more than anything else, the exact correspondence between the deterministic and stochastic models emphasizes the need for synchronized single-cell measurements: In Sec. IIF, we illustrated (i) how similarities in the effective duration of the C period obscures distinct biological mechanisms as well as (ii) how differences in the effective D period could belie an identical mechanism.

At a mechanistic level, stochasticity plays a central role in many processes. For instance, the mechanism that restarts replication will prevent the existence of a *fat tail* on the distribution of C periods [23,24,29]. Although the existence of the fat tail—*i.e*. a small number of cells with very long C periods—does not *break* the correspondence with the deterministic model, it does increase the exponential-mean C period, which in turn decreases the growth rate. (E.g. see Tab. II.) Since the growth rate is decreased, there is a strong selective pressure to reduce *stochasticity*. This argument predicts the existence of biological mechanisms to reduce stochasticity, as are already known in many contexts (*e.g*. replication restart). In fact, the subtle signature of stochasticity suggests an interesting hypothesis: a significant number of mutants that are currently known to reduce growth rate may in fact generate this phenotype by increasing the level of stochasticity in the cell cycle duration. Single-cell experimental analysis must play a central role in understanding these phenomena.

## ACKNOWLEDGMENTS

PAW acknowledges advice and comments from M. Cosentino-Lagomarsino, S. Iyer-Biswas, P. Levine, J. Mittler, R. Phillips, M. Transtrum, B. Traxler. This work was support by NIH grant GM128191.

## Appendix A: Supplemental derivations

### 1. Derivation of the solution in the deterministic model

In steady-state growth, we can assume an exponentially growing solution of the form:

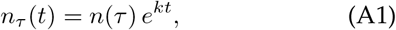

with unknown growth rate *k*. Eq. 5 can then be integrated to yield:

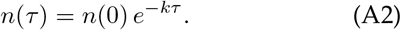

#### a. A note on normalization

Consider the creation number Eq. 12 evaluated at *τ* = 0:

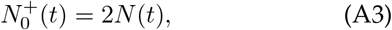

which is double the current number of cells *N*. The factor of two arises due to the cumulative nature of the creation number. To understand this intuitively, consider the total number of cells in each generation:

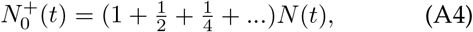

which is a geometric series and can be summed to 2*N*, matching Eq. A3.

### 2. Derivation of the solution in the stochastic model

Clearly, Eqs. 31 and 32 can be combined recursively to generate a relation between the number of cells entering states 1 and *j*. First let us define the state-transition time PDF *p_j_*, describing the total time taken to transition from birth through state *j*:

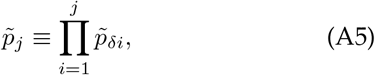

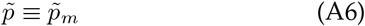

where *p* is the lifetime PDF for the entire cell cycle. We then write an expression of the number arriving in state *j*:

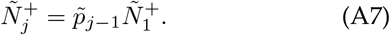

As before, using this same condition at the end of the cell cycle gives rise to a consistency condition:

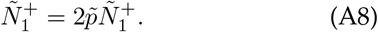

It follows that in steady-state exponential growth, the growth rate *k* must correspond to the solution to the equation:

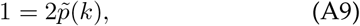

an equation that is well known [18].

Let *N*_≤*j*_ be the total number of cells in states *i* = 1…*j*. The dynamics of this quantity has a simple form due to the telescoping form of the dynamics equations (Eqs. 30-32) where the number entering the ith state exactly cancel the number leaving the *i* – 1th state:

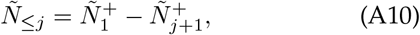

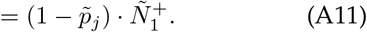

To determine the overall normalization, we can sum up the cells in all states and set that sum equal to the total number of cells *N*(*t*):

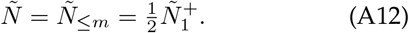

From *Ñ*_≤*j*_, we can compute the number in individual states:

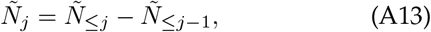

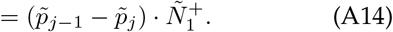

In the long time limit, the fastest growing mode dominates the solution and therefore:

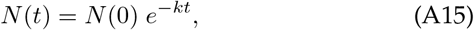

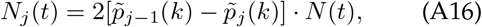

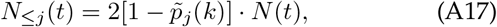

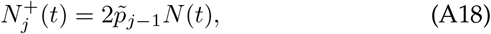

which also appear in the main text.

### 3. A generalization of the stochastic model

Like the deterministic model before it, it seems almost certain that the phenomenology of the stochastic model is more general than some of the assumptions made to motivate and derive it. In particular, the qualitative mechanism that makes the exponential-mean of state lifetime the determinative statistic would seem to depend only on the exponential enrichment of young cells in an exponential culture and not on the details of the sequential state structure of the stochastic model. We therefore offer a slightly more general derivation below.

In the generalized model, assume only that state or object *j* is created with wait time distribution *p*′ relative to the birth of a new cell and assume steady-state growth at rate *k*. Under these assumptions, 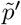 replaces 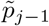 in Eqs. A7 and A18, even if *k* is not determined by Eq. A9 due to memory effects. Therefore, most of our results generalize in this new model if the suitable PDFs for the wait times replace the *p_j_*′s.

